# The *Trypanosoma brucei* Cytoskeletal Protein KHARON Associates with Partner Proteins to Mediate Both Cytokinesis and Trafficking of Flagellar Membrane Proteins

**DOI:** 10.1101/2020.10.07.330316

**Authors:** Marco A. Sanchez, Scott M. Landfear

## Abstract

In the African trypanosome *Trypanosoma brucei*, the cytoskeletal protein *Tb*KHARON is required for trafficking of a putative Ca^2+^ channel to the flagellar membrane, and it is essential for parasite viability in both the mammalian stage bloodstream forms and the tsetse fly procyclic forms. This protein is located at the base of the flagellum, in the pellicular cytoskeleton, and in the mitotic spindle in both life cycle forms, and it likely serves multiple functions for these parasites. To begin to deconvolve the functions of KHARON, we have investigated partners associated with this protein and their roles in parasite biology. One KHARON associated protein, *Tb*KHAP1, is a close interaction partner that can be crosslinked to KHARON by formaldehyde and pulled down in a molecular complex, and it colocalizes with *Tb*KHARON in the basal body at the base of the flagellum. Knockdown of *TbKHAP1* mRNA has similar phenotypes to knockdown of its partner *Tb*KHARON, impairing trafficking of the Ca^2+^ channel to the flagellar membrane and blocking cytokinesis, implying that the *Tb*KHARON/*Tb*KHAP1 complex mediates trafficking of flagellar membrane proteins. Two other KHAPs, *Tb*KHAP2 and *Tb*KHAP3, are in close proximity to *Tb*KHARON, but knockdown of their mRNAs does not affect trafficking of the Ca^2+^ channel. Two different flagellar membrane proteins, which are extruded from the flagellar membrane into extracellular vesicles, are also dependent upon *Tb*KHARON for flagellar trafficking. These studies confirm that *Tb*KHARON acts in complexes with other proteins to carry out various biological functions, and that some partners are involved in the core activity of targeting membrane proteins to the flagellum.

## INTRODUCTION

African trypanosomes of the species *Trypanosoma brucei* are parasitic protists that cause human African trypanosomiasis and the disease nagana in cattle and are thus of great medical and veterinary importance (Kennedy, 2013). In addition, these parasites have been recognized as valuable models for probing fundamental questions in cell and molecular biology (Cayla et al., 2019). *T. brucei* and related kinetoplastid parasites such as *Trypanosoma cruzi* and *Leishmania* species are flagellated and offer novel insights into the structure and function of flagella (Langousis and Hill, 2014) and the roles of these organelles in infection (Kelly et al., 2020a). Furthermore, the cell division cycle of African trypanosomes has been studied extensively (Farr and Gull, 2012; Vaughan and Gull, 2008; Wheeler et al., 2019), identifying various processes that are important for both proliferative and differentiation-linked cell division.

In previous work on trafficking of integral membrane proteins to flagella, we identified a kinetoplastid-specific protein designated KHARON (KH) that plays a critical role in flagellar targeting of the putative Ca^2+^ channel *Tb*CaCh (Tb927.10.2880) in *T. brucei*, originally identified as a flagellar surface protein FS179 (Oberholzer et al., 2011), and the flagellar glucose transporter *Lmx*GT1 in *L. mexicana* (Tran et al., 2013) (see Table S1 for tabulation of gene IDs and names of *T. brucei* proteins investigated in this study). In both species of parasite (Sanchez et al., 2016), KHARON was localized to three distinct subcellular compartments: the base of the flagellum (Fig. 1A), the subpellicular microtubules that subtend the plasma membrane around the cell body, and the mitotic spindle (Fig. 1B). Application of RNA interference (RNAi) to knock down *TbKH* (Tb927.10.8940) revealed that, in addition to preventing flagellar trafficking of *Tb*CaCh/FS179, the flagellum attachment zone (FAZ) was disrupted, resulting in detachment of flagella from the cell body (Sanchez et al., 2016), leaving this organelle adherent only through its connection at the flagellar pocket. Interfering with trafficking of *Tb*CaCh/FS179 to the flagellar membrane is likely to induce disruption of flagellar attachment, as RNAi directed against this channel also results in a similar flagellar detachment phenotype (Oberholzer et al., 2011). In addition, as found for many genetic alterations that disrupt flagellar attachment, these parasites are unable to initiate cell division, generating trypanosomes in which nuclei, kinetoplasts (mitochondrial DNA-containing structures), basal bodies, and flagella have replicated but cytokinesis has not occurred. This phenotype was apparent in both mammalian bloodstream form (BF) and insect stage procyclic form (PF) parasites and was thus lethal to both life cycle stages.

**Fig. 1.**
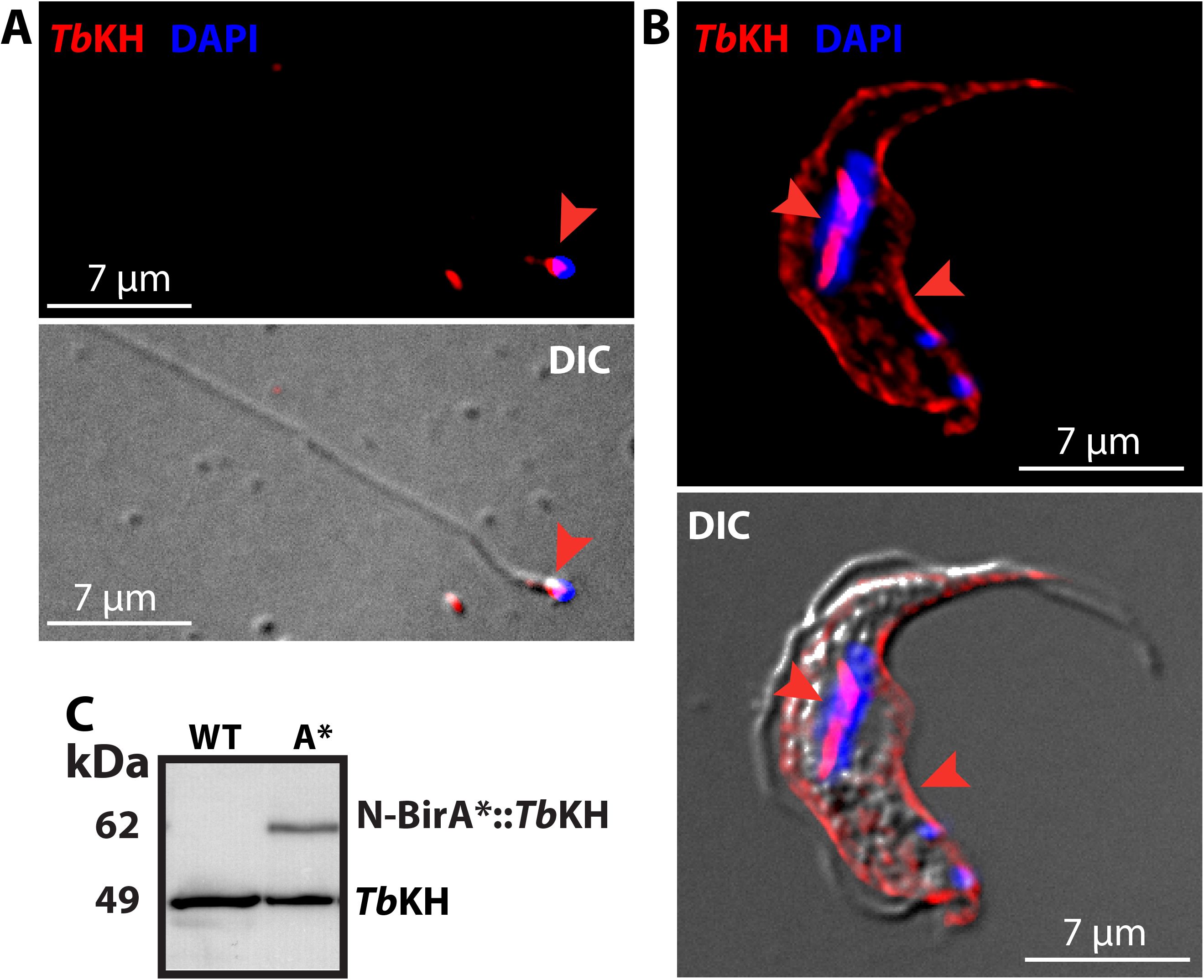
*Tb*KH localizes in three distinct subcellular compartments. (A) Flagellar preparation from *Tb*FLA1^RNAi^ cell line, isolated after 72 h of induction with 1 μg/ml doxycycline (dox), stained with DAPI (*blue*) and immunostained with anti-*Tb*KH pAb (*Tb*KH, *red*). *Tb*KH localization at the base of the flagellum is indicated by the *red arrowhead*. (B) Wild type BF parasites were immunostained with anti-*Tb*KH pAb (*Tb*KH, *red*) and stained with DAPI. These images display a parasite in which *Tb*KH staining is associated with both the subpellicular microtubules (*right arrowheads*) and the mitotic spindle that connects the two nuclei late during mitosis (*left arrowheads*). DIC indicates images collected by differential interference contrast microscopy. (C) Western blot of total protein lysates from BF trypanosomes that are either wild type (*WT*) or expressing BirA*::TbKH fusion protein (*A\**). Blot was probed with anti-*Tb*KH pAb and developed by chemiluminescence. Specific immunodetected proteins corresponding to the native *Tb*KH and the BirA*::*Tb*KH fusion protein are shown. Relative protein molecular weights are shown in kDa, as determined by mobility relative to molecular weight markers.

**Table 1.** List of Proteins with Gene IDs.

| Gene ID | Protein |
| --- | --- |
| Tb927.10.8940 | *Tb*KH |
| Tb927.10.2880 | *Tb*CaCh/FS179 |
| Tb927.11.2610 | *Tb*KHAP1 |
| Tb927.10.10360 | *Tb*KHAP2/MARP-1 |
| Tb927.10.10280 | *Tb*KHAP3/MARP-2 |
| Tb927.7.4270 | *Tb*EVMP1/Fam79.3 |
| Tb927.11.1830 | *Tb*EVMP2 |
| Tb927.8.4050 | *Tb*FLA1BP |

Similar studies in *L. mexicana* have also established a critical role for *Lmx*KH (LmxM.36.5850) in the life cycle of *Leishmania* parasites. Thus a Δ*lmxkh* null mutant was generated in insect stage promastigotes, where trafficking of *Lmx*GT1 to the flagellum was strongly impaired (Tran et al., 2013), but cell division and replication of this life cycle stage was not affected. In contrast, Δ*lmxkh* null mutants were unable to undergo cytokinesis after invading host macrophages, resulting in formation of multinucleate multiflagellated amastigotes that died over the course of several days (Tran et al., 2013). These null mutants were also avirulent following injection into BALB/c mice (Tran et al., 2015), and studies by others have shown that *KHARON* null mutants in *L. infantum* have potential as a live attenuated vaccine (Santi et al., 2018).

KHARON exhibits both similarities and striking differences between *T. brucei* (*Tb*KH) and *L. mexicana* (*Lmx*KH). Thus, the two orthologs are relatively divergent in sequence, sharing 27% amino acid identity and differing significantly in length (411 amino acids for *Tb*KH versus 520 amino acids for *Lmx*KH). *Tb*KH is critical for cell division of both mammalian BF and insect stage PF parasites, whereas *Lmx*KH is only essential for division of disease-causing amastigotes. Nonetheless, the three subcellular locations for KHARON are shared between the two parasites, as are functions in cytokinesis and formation of the flagellar membrane.

KHARON proteins do not share significant sequence similarity to proteins outside the Kinetoplastida, nor do they contain conserved sequence motifs that are suggestive of specific biochemical or cellular functions. Furthermore, their residence at multiple subcellular locations suggests that KHARON proteins are likely to be multifunctional, participating in flagellar membrane trafficking, cytokinesis, and spindle function. Additionally, it is possible that distinctions in function could be conferred by association of KHARON with different partner proteins at each of its three subcellular loci. Thus, we hypothesize that there could exist three distinct KHARON Complexes, Complex 1 at the base of the flagellum, Complex 2 at the subpellicular cytoskeleton, and Complex 3 at the mitotic spindle. Furthermore, given the apparent differences in *Tb*KH and *Lmx*KH noted above, there may be similarities and differences between these putative complexes between the two species of parasite.

To initiate a study of putative KHARON Complexes and their functions, we carried out biotinylation proximity labeling (BioID) (Roux et al., 2012) and tandem affinity purification-mass spectrometry (TAP-MS) (Kaiser et al., 2008) on *Lmx*KH, resulting in the identification of two KHARON Associated Proteins, *Lmx*KHAP1 and *Lmx*KHAP2 (LmxM.32.2440 and LmxM.05.0380, respectively; (Kelly et al., 2020b)). In parallel, we investigated these two KHARON partners in *T. brucei* and report the results of those studies here. As anticipated, *Tb*KHAP1 and *Tb*KHAP2 exhibit both similarities and notable differences compared to their orthologs in *L. mexicana*. In addition, *T. brucei* expresses another KHARON partner related to *Tb*KHAP2 that we designate *Tb*KHAP3. Furthermore, characterization of additional flagellar membrane proteins suggests that *Tb*KH expression is important for flagellar targeting of multiple such proteins in African trypanosomes, whereas the role of *Lmx*KH in trafficking of flagellar membrane proteins appears to be more restricted. These studies confirm that KHARON proteins in both parasites exist in complexes with various partners and that these partner proteins can play distinct roles in the functions of different KHARON Complexes.

## RESULTS

### Localization of *Tb*KHAP1, *Tb*KHAP2, TbKHAP3, and *Tb*KH in bloodstream and procyclic African trypanosomes

To facilitate studies on *Tb*KH and its partners, we raised and affinity purified a polyclonal antibody against this protein, anti-*T*bKH pAb. Western blot analysis indicated that anti-*Tb*KH pAb detects a single protein of ~49 kDa molecular weight, and that an additional band of ~62 kDa appears in parasites also expressing a BirA* fusion on the N-terminus of *T*bKH (Fig. 1 C), establishing that this antibody is of suitable specificity to employ in localization and biochemical characterization of *Tb*KH. To determine whether *Tb*KHAP1 (Tb927.11.2610), *Tb*KHAP2 (Tb927.10.10360) and *Tb*KHAP3 (Tb927.10.10280) are associated with *Tb*KH in a complex, several complementary approaches were applied. First, each KHAP was tagged at its N-terminus with the triple hemagglutinin peptide tag HA_3_, and formaldehyde-fixed BF and PF trypanosomes were examined by immunofluorescence deconvolution microscopy (Fig. 2). HA_3_::*Tb*KHAP1 (Fig. 2A,B, green) overlaps with *Tb*KH (red) at the cell periphery, as demonstrated by the yellow color in this region of both BF and PF parasites. In contrast, there was no apparent overlap of the two signals in the mitotic spindle (central red oval or line marked with a white arrow), indicating that this protein could be associated with *Tb*KH in the subpellicular cytoskeleton but not at the mitotic spindle. Similarly, *Tb*KHAP2::HA_3_ (Figs. 2C,D) and V5_3_::*Tb*KHAP3 or HA_3_::*Tb*KHAP3 (Fig. 2E,F) overlap with *Tb*KH at the cell periphery but not at the mitotic spindle. For each of the *Tb*KHAPs, there is also green fluorescence that does not coincide with *Tb*KH so that there is not complete overlap of the signals, and there may thus be populations of each protein that are not associated with each other. However overall, these three *Tb*KHAPs are candidates for *Tb*KH partners that are selective for the subpellicular cytoskeleton versus the mitotic spindle.

**Fig. 2.**
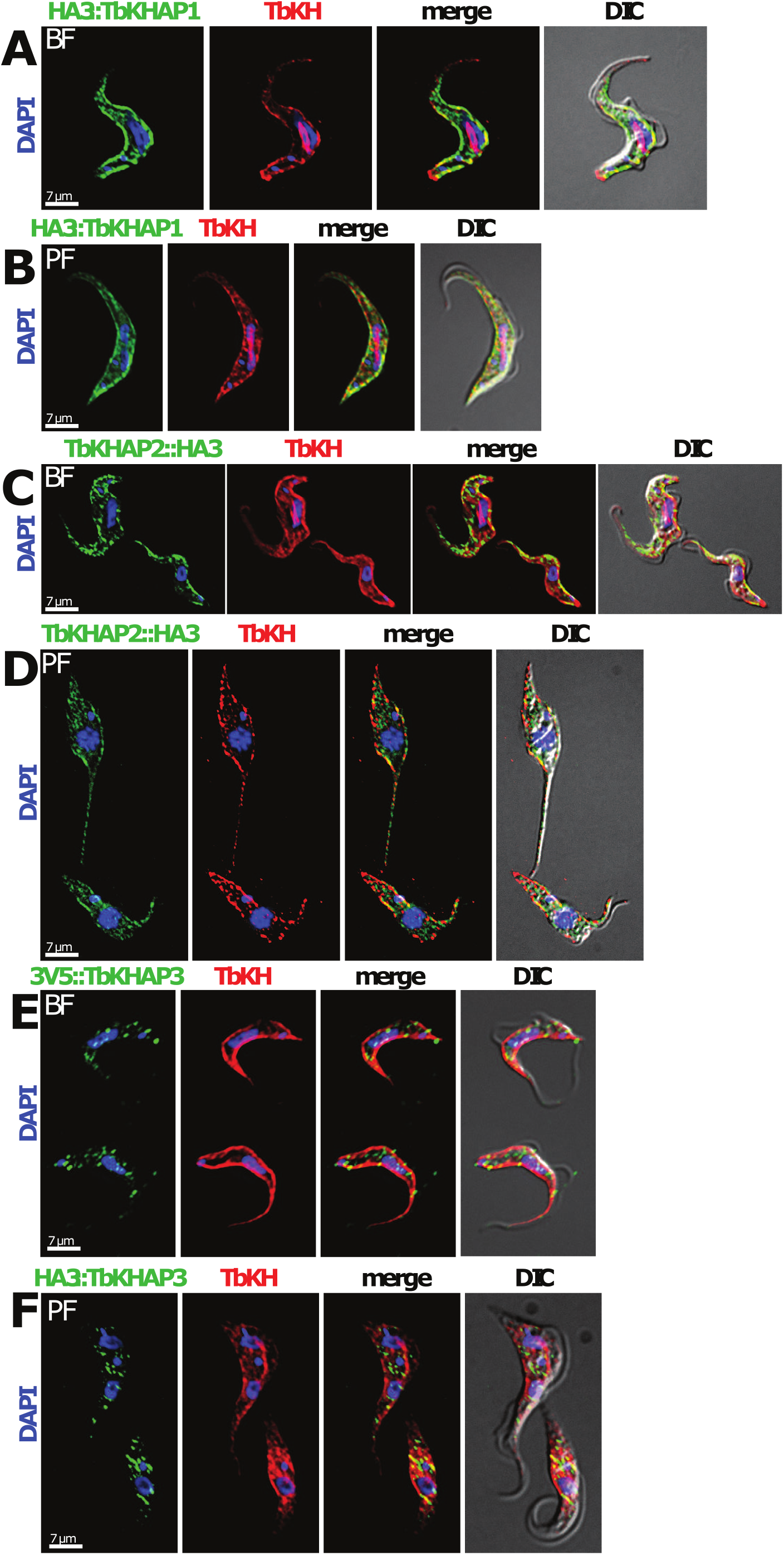
Subcellular localization of *Tb*KHAPs in BF and PF *T. brucei*. (A, B) The HA_3_::TbKHAP1 cell line was immunostained with anti-HA mAb (HA_3_::*Tb*KHAP1*, green*) and anti-*Tb*KH pAb (*Tb*KH*, red*). (C, D) The TbKHAP2::HA_3_ cell line was stained with anti-HA mAb (*Tb*KHAP2::HA_3_*, green*) and anti-*Tb*KH pAb (*Tb*KH*, red*). (E) The BF 3V5::*Tb*KHAP3 and (F) PF HA_3_::*Tb*KHAP3 cell lines were immunostained with (E) anti-V5 mAb (3V5::*Tb*KHAP3, *green)* or (F) anti-HA mAb (HA_3_::*Tb*KHAP3*, green*) and anti-*Tb*KH pAb (*Tb*KH*, red*). All preparations were stained with DAPI, which detects both nuclear and kinetoplast DNA (*blue*). DIC images were also acquired from all samples.

To determine whether *Tb*KHAP1, *Tb*KHAP2, or *Tb*KHAP3 might associate with *Tb*KH at the base of the flagellum, flagella were isolated from parasites expressing each HA_3_-tagged *Tb*KHAP and imaged by deconvolution microscopy. Fig. 3 shows that HA_3_::*Tb*KHAP1 (Fig. 3A), HA_3::_TbKHAP2 (Fig. 3B) and HA_3_::TbKHAP3 (Fig. 3C) overlap significantly with *Tb*KH in the region of the flagellum immediately adjacent to the kinetoplast DNA (kDNA, blue, Fig. 3A), which is close to and physically attached to (Robinson and Gull, 1991) the flagellar basal body. Additional images showing overlap of *Tb*KH with each of the three TbKHAPs at the base of the flagellum are provided in Fig. S1. Overall, these results indicate that *Tb*KHAP1, *Tb*KHAP2, and *Tb*KHAP3 could be partners for *Tb*KH at both the pellicular cytoskeleton and the base of the flagellum. It is noteworthy that *Tb*KHAP3 was previously identified as a basal body protein in a proteomic study of that subcellular structure (Dang et al., 2017). Hence, it is likely that *Tb*KH, *Tb*KHAP1, *Tb*KHAP2, and *Tb*KHAP3 are all basal body components.

**Fig. 3.**
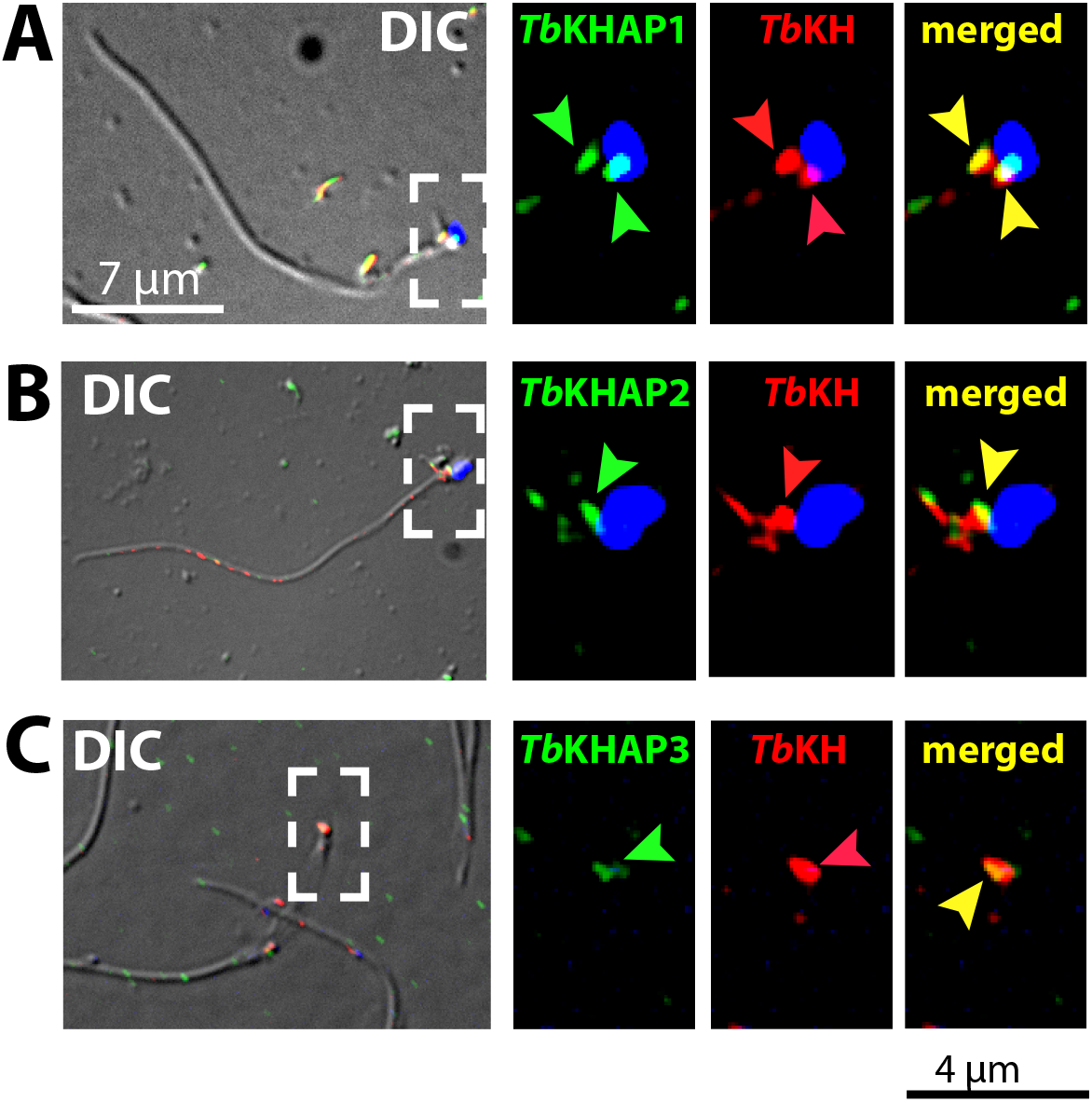
KHARON and KHAPs colocalize at the base of the flagellum in *T. brucei*. (A) Whole isolated flagella from the BF HA_3_::*Tb*KHAP1/*Tb*FLA1^RNAi^ clone induced for RNAi for 72 h, were immunostained with anti-HA mAb (HA_3_::*Tb*KHAP1, *green*) and anti-*Tb*KH pAb (*Tb*KH, *red*) and stained with DAPI (*blue*). (A, left panel) shows an immunofluorescence/DIC image of an isolated flagellum, *white punctuated* box indicates the magnified area depicted in (A, center-right, center-left and left) panels. *Green arrowheads* indicate HA_3_::*Tb*KHAP1 localization, *red arrowheads* indicate *Tb*KH localization and *yellow arrowheads* indicate overlapping signals, when channels are merged, near the kinetoplast (*blue*). (B) Flagella were isolated from BF trypanosomes expressing HA_3_::TbKHAP2 and imaged as described in (A). (C) Flagellar cytoskeletons were isolated from PF trypanosomes expressing HA_3_::TbKHAP3 and imaged as described in (A).

### Molecular association of *Tb*KHAP1, *Tb*KHAP2, and *Tb*KHAP3 with *Tb*KH

To determine whether the observed subcellular overlap of the fluorescence signals from *Tb*KHAPs and *Tb*KH could indicate physical association in molecular complexes, *Tb*KH was endogenously tagged at its N-terminus with a His_10_ affinity tag (His_10_::*Tb*KH) to allow pulldown of this protein, and associated partners, with Ni-NTA magnetic beads, and this affinity tagged protein was expressed in a BF cell line also expressing either HA_3_::*Tb*KHAP1, *Tb*KHAP2:: HA_3_, or V5_3_::*Tb*KHAP3. Because *Tb*KH is an integral component of the parasite cytoskeleton (Sanchez et al., 2016) and would pulldown many cytoskeletal proteins in an experiment performed under native conditions, we first crosslinked with formaldehyde parasites expressing each pair of tagged proteins. Formaldehyde crosslinks proteins that are in very close proximity (~2-3 Å, reference (Hoffman et al., 2015)), so this treatment will covalently attach close molecular partners of *Tb*KH but not proteins that are more distant partners in a complex or proteins that are in the cytoskeleton but distant from *Tb*KH. Subsequent treatment with strongly denaturing reagents will dissociate peripheral proteins from His_10_::*Tb*KH while retaining crosslinked partners, and the closely associated partners will thus be purified along with His_10_::*Tb*KH, following binding and elution from the Ni-NTA beads, and released upon heat-induced reversal of the crosslinks.

Fig. 4A demonstrates that HA_3_::*Tb*KHAP1 is pulled down with His_10_::*Tb*KH when parasites are formaldehyde crosslinked (EF*) but not when they are not subjected to crosslinking (EF). As a negative control, another subpellicular cytoskeletal protein, CAP15 (Vedrenne et al., 2002), was HA_3_ tagged and expressed in His_10_::*Tb*KH expressing BF parasites, but this protein was not pulled down even from formaldehyde-crosslinked parasites. Parallel experiments were performed using BF parasites expressing *Tb*KHAP2::HA_3_, or V5_3_::*Tb*KHAP3 and His_10_::*Tb*KH. However, for reasons that are not clear but possibly having to do with the highly repetitive nature of both proteins (see below), treatment of parasites with formaldehyde followed by immediate dissolution in the strongly denaturing urea buffer employed to disrupt protein interactions led to massive degradation of both TbKHAP2::HA_3_ and TbKHAP3::HA_3_. Indeed, the susceptibility of these proteins to degradation has been noted previously (Schneider et al., 1988). Hence, for technical reasons we have not been able to observe pulldowns of either of these two tagged protein with His_10_::*Tb*KH, thus preventing us from directly demonstrating interactions between these proteins at the 2-3 Å level. Nonetheless, these experiments establish that *Tb*KHAP1 is in very close proximity to *Tb*KH.

**Fig. 4.**
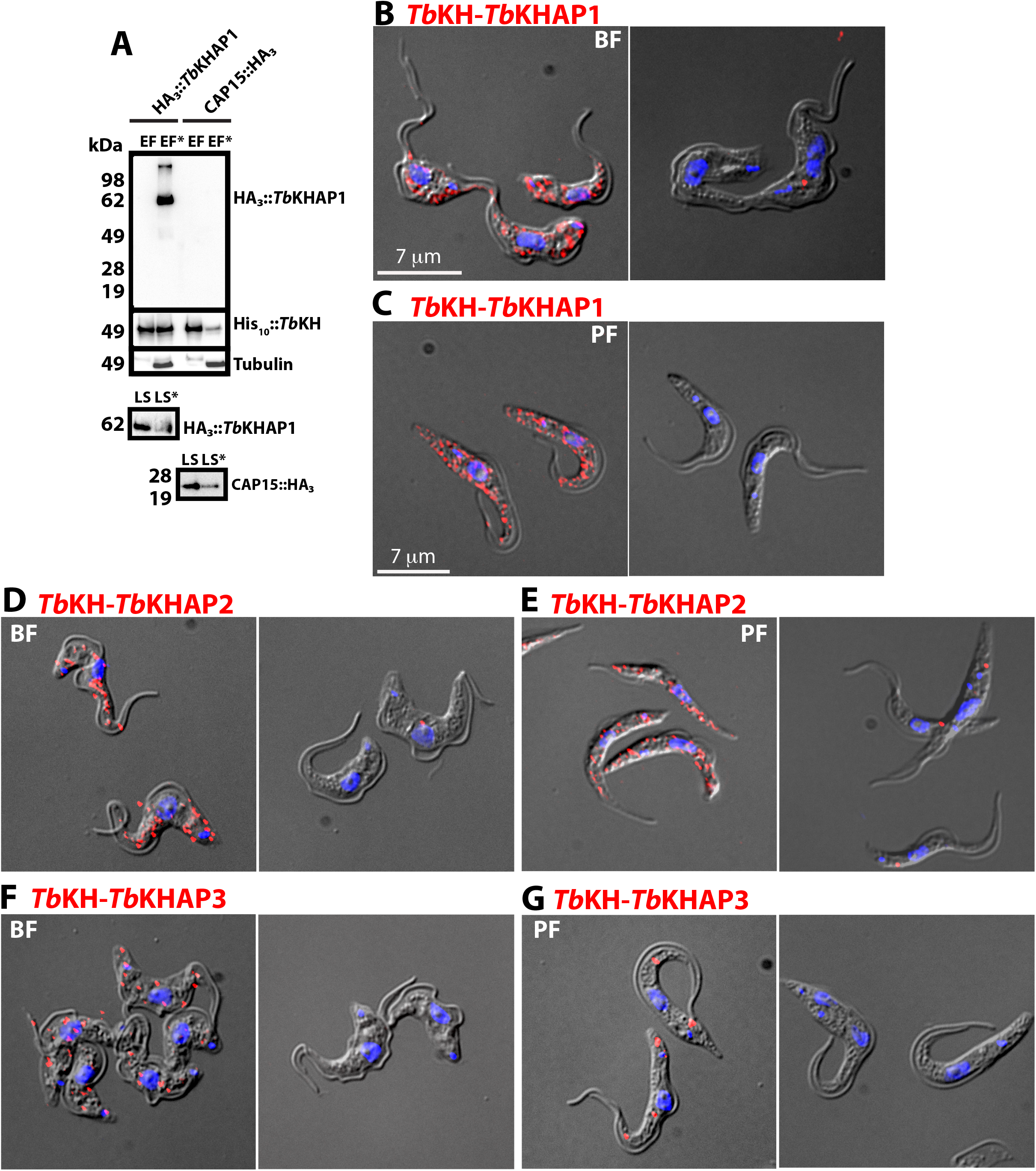
*Tb*KH is closely associated with *Tb*KHAP1, *Tb*KHAP2, and *Tb*KHAP3. (A) Western blot of protein samples from pull down of HA_3_::*Tb*KHAP1 using His_10_::*Tb*KH as a bait and CAP15::HA_3_ as negative control. *LS* and *EF*, lysate and elution fractions without formaldehyde crosslinking; *LS\** and *EF*\*, lysate and elution fraction with formaldehyde crosslinking. Protein blots were developed by chemiluminescence, bands are indicated by protein names, and migration of molecular weight markers are designated in kDa. (B, C) PLA employing BF or PF HA_3_::TbKHAP1/His_10_::TbKH cell lines, as indicated. For these and all subsequent sections of this figure, the *left panel* shows results using both anti-HA mAb and anti-*Tb*KH pAb, and the *right panel* shows results using only the anti-*Tb*KH pAb, as the negative control. *Red puncta* indicate positive HA_3_::*Tb*KHAP1-*Tb*KH interaction. DIC images were acquired from all samples. (D, E) PLA employing BF or PF *Tb*KHAP2::HA_3_ cell lines, as indicated. *Red puncta* indicate positive *Tb*KHAP2-*Tb*KH interaction. (F, G) PLA employing BF or PF HA_*3*_::*Tb*KHAP3 cell lines, as indicated using. *Red puncta* indicate positive *Tb*KHAP3-*Tb*KH interaction.

### Proximity ligation assay (PLA) confirms close proximity of *Tb*KH with *Tb*KHAP1, *Tb*KHAP2, and *Tb*KHAP3

To provide another independent examination of whether *Tb*KH is in close physical proximity with each *Tb*KHAP, we performed the PLA in parasites expressing HA_3_::*Tb*KHAP1, *Tb*KHAP2::HA_3_, and HA_3_::*Tb*KHAP3. In this assay (Fredriksson et al., 2002; Soderberg et al., 2006), cells expressing two partner proteins are first reacted with primary antibodies from different species. Parasites are subsequently incubated with species-specific secondary antibodies directed against each primary antibody, and each of these secondary antibodies contains a unique, covalently attached oligonucleotide. Only if the two target proteins are within ~400 Å of each other, these oligonucleotides can base pair to another linker oligonucleotide and be covalently ligated into a circular substrate that can participate in rolling circle DNA amplification of the cognate sequence. The amplified sequence is then hybridized to a fluorescently labeled DNA probe, resulting in fluorescent puncta within the cell.

Fig. 4B,C shows a positive PLA signal (left panels) for BF and PF trypanosomes expressing HA_3_::*Tb*KHAP1 and probed with anti-*Tb*KH rabbit and anti-HA murine mAb. In contrast, when the anti-HA murine antibody is not employed (right panels), the PLA signal is absent, demonstrating the dependency of the signal on detection of both closely associated proteins. Similar results confirm that *Tb*KH is in close physical proximity to *Tb*KHAP2 (Fig. 4D,E) and *Tb*KHAP3 (Fig. 4F,G).

### Predicted properties of *Tb*KHAP1, *Tb*KHAP2, and *Tb*KHAP3

Bioinformatic analysis of the 50.9 kDa *Tb*KHAP1 sequence indicates that it is a protein apparently unique to kinetoplastid protists for which there are orthologs widely distributed among Kinetoplastida. A BLASTP search (tritrypdb.org) revealed several coiled-coil proteins such as neurofilament proteins and tropomyosin as being significantly, although not closely, related (E values of 2.6e-08 – 5.4e-12). Prediction of protein disorder using the PrDOS web server (Ishida and Kinoshita, 2007) (http://prdos.hgc.jp/cgi-bin/top.cgi) generated a strong prediction of disorder (probability >0.9) over the C-terminal region from amino acids 314 – 461. Indeed, this sequence is rich in E residues, which predispose such regions to intrinsic disorder (Uversky, 2013), and this property suggests that this region could be involved in protein-protein interactions through induced folding (Zhang et al., 2013). InterPro (Mitchell et al., 2015) (https://www.ebi.ac.uk/interpro/search/sequence/) predicted coils between amino acids 7 – 31 and 196 – 241 and a disordered region from residue 332 - 461, and PSIPRED V4.0 (McGuffin et al., 2000) (http://bioinf.cs.ucl.ac.uk/psipred/) predicted the sequence to be largely helix or coil. Overall, computational analyses suggest that TbKHAP1 is a coiled-coil protein with an intrinsically disordered C-terminus, both properties that are consistent with formation of multi-protein complexes.

*Tb*KHAP2 is the 374 kDa microtubule-associated repetitive protein 1, MARP-1, and *Tb*KHAP3 is the 267 kDa MARP-2 that have been studied previously by Seebeck and colleagues (Affolter et al., 1994; Hemphill et al., 1992; Schneider et al., 1988) and will hereafter be referred to as *Tb*KHAP2/MARP-1 and *Tb*KHAP3/MARP-2 to indicate both their association with *Tb*KH and their previously demonstrated roles in microtubule binding. Each sequence contains short unique N- and C-terminal domains, and the remainder of the sequence consists of 38-amino acid repeats that are largely conserved within each sequence but ~50% identical between the two proteins. The unique C-terminal domains (95% identical between the two proteins) bind to microtubules (Affolter et al., 1994), and the proteins decorate the subpellicular cytoskeleton (Schneider et al., 1988), but their specific biological functions have not been elucidated. In addition, these proteins have also been localized to the basal body and *Tb*KHAP3/MARP-2 was designated *Tb*BBP268 (Dang et al., 2017).

### Phenotypes of BF trypanosomes following knockdown of *TbKHAP1* RNA

We have previously demonstrated that knockdown of *TbKH* RNA by inducible RNAi results in a lethal phenotype on both BF and PF trypanosomes (Sanchez et al., 2016). In these parasites, the flagellum detaches from the cell body along the flagellum attachment zone (FAZ), and the parasites are blocked in cytokinesis, resulting in accumulation of multi-nucleated, multi-flagellated ‘monster cells’ that are not viable in the long term. To assess the roles of *Tb*KHAP1, *Tb*KHAP2/MARP-1, and *Tb*KHAP3/MARP-2 in the biology of BF parasites, we targeted by RNAi *TbKHAP1* mRNA, using a unique RNAi probe, and *TbKHAP2/TbKHAP3* mRNAs jointly, using a 500 nt probe covering the conserved C-termini, and assessed the consequent phenotypes.

Induction of RNAi against *Tb*KHAP1 in BFs using doxycycline resulted in rapid reduction in the level of this protein (Fig. 5A). Furthermore, RNAi-induced parasites stopped growing almost immediately and were largely dead by 72 h (Fig. 5B). Quantification via microscopy of the percentage of cells with different numbers of nuclei and kinetoplasts (Fig. 5C) showed that following induction of RNAi over 48 h, the percentage of 1N1K parasites dropped dramatically, while those with multiple nuclei and kinetoplasts (XNYK) increased and began to predominate the population. In comparison to the normal morphology of pre-induced parasites (Fig. 5D), induction of RNAi for 20 h (Fig. 5E) or 48 h (Fig. 5F) resulted in parasites with multiple nuclei and/or kinetoplasts and tadpole-like morphology (white arrowhead) or duplicated flagella located at opposite poles of the cell body (yellow arrowhead).

**Figure 5.**
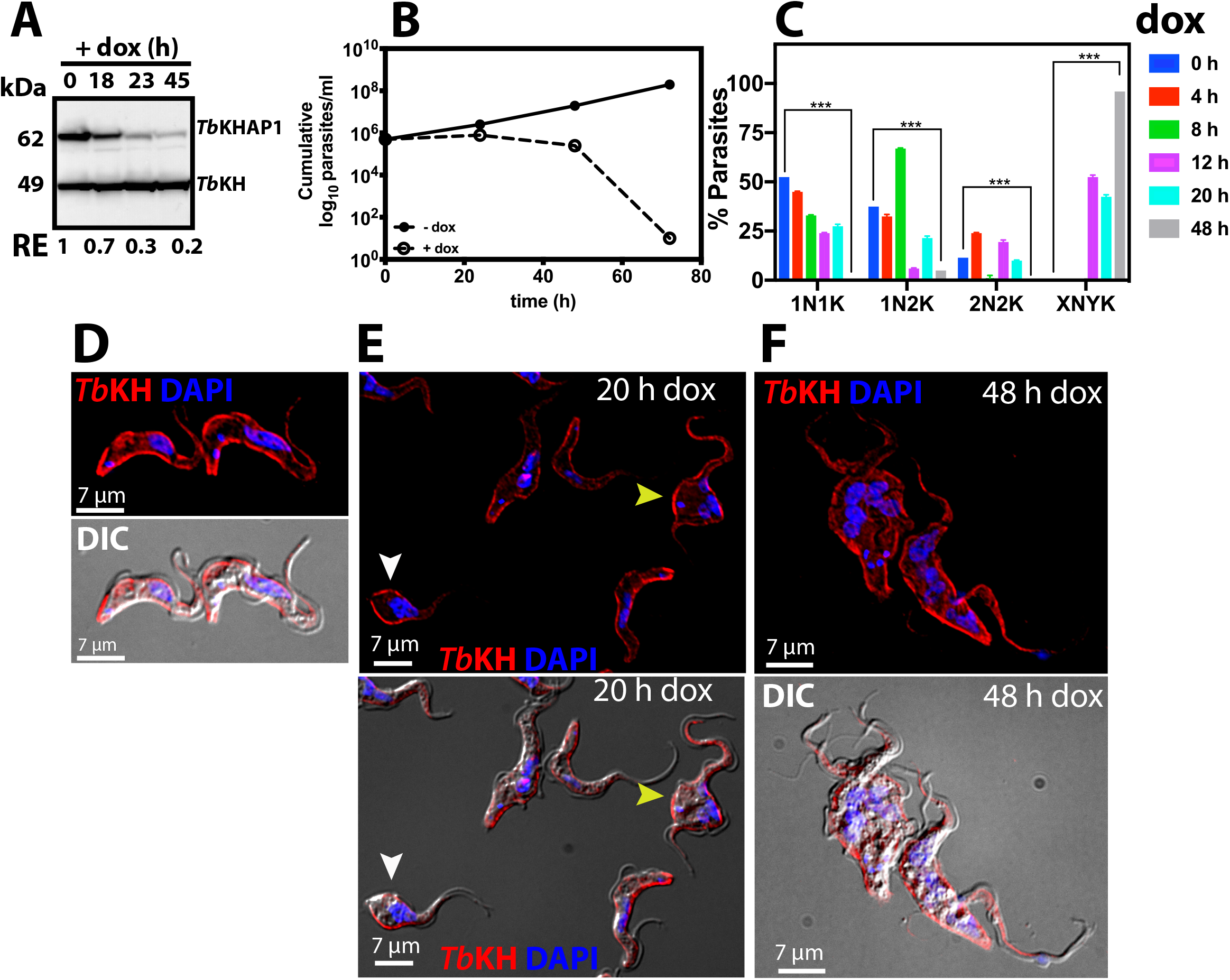
Depletion of *Tb*KHAP1 is lethal for *T. brucei* parasites. (A) Western blot of total protein lysates from TbKHAP1^RNAi^ BF parasites grown in the presence of doxycycline (+dox) and immunodetected with anti-HA mAb, anti-*Tb*KH pAb. The approximate molecular weights in kDa of HA_3_::*Tb*KHAP1 and *Tb*KH are indicated as determined by mobility compared to weight markers. *Numbers* under the blot represent the relative intensity (*RI*) of HA_3_::*Tb*KHAP1 protein normalized to the *Tb*KH protein level. (B) Cell density for induced (*empty circles*) and non-induced (*filled circles*) *Tb*KHAP1^RNAi^ BF cell lines. Parasite density was quantified by phase contrast microscopy using a hemocytometer. Data represent the averages and standard deviations of two experiments, each employing an independently isolated *Tb*KHAP1^RNAi^ clonal cell line, and technical replicates were also performed for each biological replicate. Standard deviations are too small to be visible. (C) Microscopic analysis of nuclei and kinetoplast numbers of BF *Tb*KHAP1^RNAi^ parasites following induction of RNAi. Data represent frequency (*%*) of cells with different numbers of DAPI stained nuclei (*N*) and kinetoplasts (*K*). Results represent the average and range of two independent experiments. (D-F) *Tb*KHAP1^RNAi^ BF parasites were stained with DAPI (*blue*) and immunostained with anti-α-tubulin mAb (Tub, *red*) at (D) 0 h, (E) 20 h and (F) 48 h post-RNAi induction. In (E) the *white arrowhead* indicates a cell with a tadpole-like morphology and the *yellow arrowhead* indicates a cell with two flagella at opposite poles of the cell body.

Notable among cells in RNAi induced populations are those with multiple flagella located at various relative positions around the cell (e.g., the two parasites in Fig. 5F). Such parasites have initiated cytokinesis and cleavage furrow formation, as the two duplicated flagella have moved apart from the initial position they would have following flagellar duplication. However, the cleavage furrow did not progress to separate the duplicated nuclei and kinetoplasts as it would in normal cell division. In summary, these observations suggest that loss of *Tb*KHAP1 protein from BF parasites results in a block in progression of cleavage furrow rather than an inability to initiate cleavage furrow ingression.

These results are further enhanced by more refined time course studies shown in Fig. S2 following the progression of nuclear content and cell morphologies between 0 – 48 h after induction of RNAi against *TbKHAP1* RNA. At 4 h (Fig. S2B) and 8 h (Fig. S2C) most parasites had morphologies similar to that preceding induction of RNAi (0 h, Fig. S2A). However, by 12 h (Fig. S2D), multi-flagellated parasites with ingression furrows appeared, and by 24 h and 48 h (Fig. S2E,F), many parasites had incompletely resolved ingression furrows and multiple nuclei.

### Phenotypes of parasites following knockdown of *TbKHAP2/MARP-1* and *TbKHAP3/MARP-2* RNAs

Induction of RNAi jointly against *TbKHAP2/MARP-1* and *TbKHAP3/MARP-2* resulted in complete loss of HA_3_::*Tb*KHAP2/MARP-1 protein by 24 h (Fig. 6A. Depletion of *Tb*KHAP2/MARP-1 and *Tb*KHAP3/MARP-2 impaired growth of BF parasites, resulting in an ~50-fold reduction in parasite number by 120 h (Fig. 6B) post-induction, but growth inhibition was not nearly as pronounced as it is for *Tb*KHAP1 RNAi (Fig. 5B). Compared to uninduced parasites (Fig. 6C), images of parasites following 4 d RNAi (Fig. 6D) still showed many parasites with normal morphology similar to that of uninduced parasites, but some parasites rounded up and showed multiple flagella (Fig. 6D, white arrowhead, flagella on opposite sides of the cell body in DIC image). By 4 d post-RNAi, parasites with 1N2K and XNYK began to accumulate (Fig. 6E), but the proportion was not nearly as great as for RNAi directed against TbKHAP1 (Fig. 5C).

**Fig. 6.**
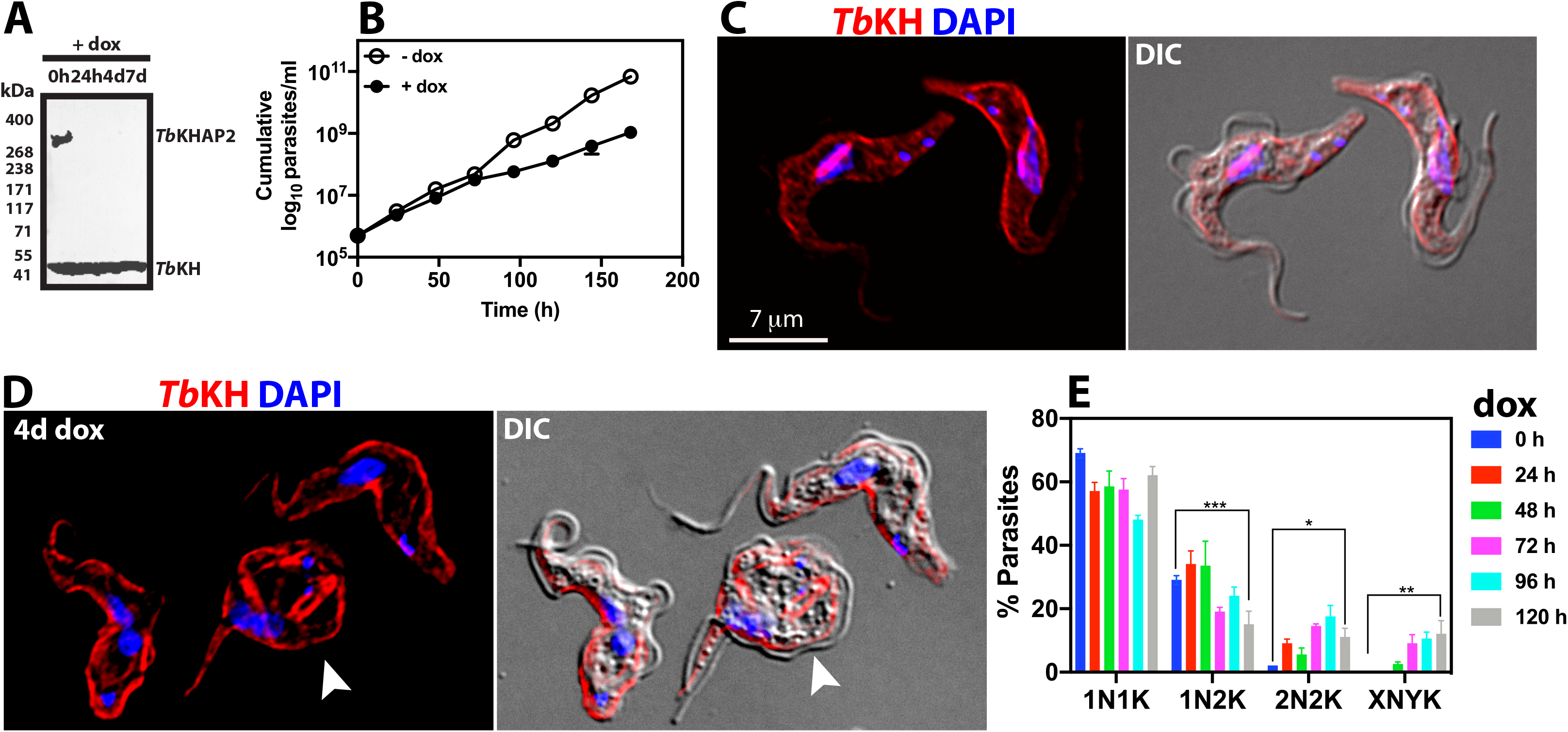
Phenotypes resulting from RNAi directed against *TbKHAP2* and *TbKHAP3*. (*A*) Western blot of total protein lysates of BF parasites expressing HA_3::_*Tb*KHAP2 following induction of RNAi (1 μg ml^−1^ doxycycline, *dox*) directed against *TbKHAP2/3* RNAs. Protein blots were probed with anti-HA mAb and anti-*Tb*KH pAb as loading control. Molecular weight markers are indicated in kDa. (*B*) Growth curve of induced (*empty circles*) and non-induced (*filled circles*) TbKHAP2/3^RNAi^ cell line. Parasite density was quantified by phase contrast microscopy using a hemocytometer. The data represent two biological replicate experiments, but the standard deviations are too small to see in the figure. Representative *Tb*KHAP2/3^RNAi^ cells induced for RNAi were stained with DAPI (*blue*) and immunostained with anti-*Tb*KH pAb (*Tb*KH, *red*) at (*C)* 0 h and (*D)* 4 days post-RNAi induction. *White arrowhead* in (*D*) indicates a cell showing aberrant morphology. DIC images were obtained for all samples. (*E*) Microscopic analysis of NK in BF parasites undergoing RNAi. Frequency (*%*) of cells with different numbers of nuclei (*N*) and kinetoplasts (*K*) at different times following induction of RNAi against *TbKHAP2/3* mRNA. Results represent the average and range of two independent experiments.

### *Tb*KHAP1, but not *Tb*KHAP2/MARP-1 or *Tb*KHAP3/MARP-2, is required for targeting *Tb*CaCh/FS179 to the flagellar membrane

The observation that *Tb*KHAP1 is located at the base of the flagellum (Fig. 3A), likely in the basal body, raises the question of whether it could play a role in the function of *Tb*KH in mediating trafficking of the putative Ca^2+^ channel, *Tb*CaCh/FS179, to the flagellar membrane. To test this possibility, we induced RNAi against *TbKHAP1* RNA in parasites expressing *Tb*CaCh/FS179::HA_3_ tagged at the C-terminus, which localized to the flagellar membrane prior to RNAi (Fig. 7A). BF parasites induced for RNAi for 24 h (Fig. 7B) or 48 h (Fig. 7C) exhibited flagella that were devoid of *Tb*CaCh::HA_3_ (white arrowheads). These results suggest that a complex of *Tb*KH/*Tb*KHAP1, and potentially other currently unknown partners located at the base of the flagellum, is involved in trafficking this channel to the flagellar membrane. Since both proteins are also located in the subpellicular cytoskeleton, it is not possible to definitively ascribe this flagellar trafficking phenotype to the complex at the base of the flagellum; complexes at both locations will be downregulated by *TbKHAP1* RNAi. However, integral membrane proteins are first delivered to the flagellar pocket membrane during biosynthesis (Manna et al., 2014). Hence, the presence of a protein complex located close to the interface between the flagellar pocket and flagellar membrane, and for which downregulation of both known partners inhibits trafficking of a protein into the flagellar membrane, suggests that this complex may mediate trafficking of *Tb*CaCh/FS179 from the flagellar pocket membrane into the flagellar membrane. Such trafficking would presumably be mediated by a direct interaction between *Tb*KH and the cargo, *Tb*CaCh/FS179, and such a molecular interaction has been demonstrated to occur (Sanchez et al., 2016).

**Fig. 7.**
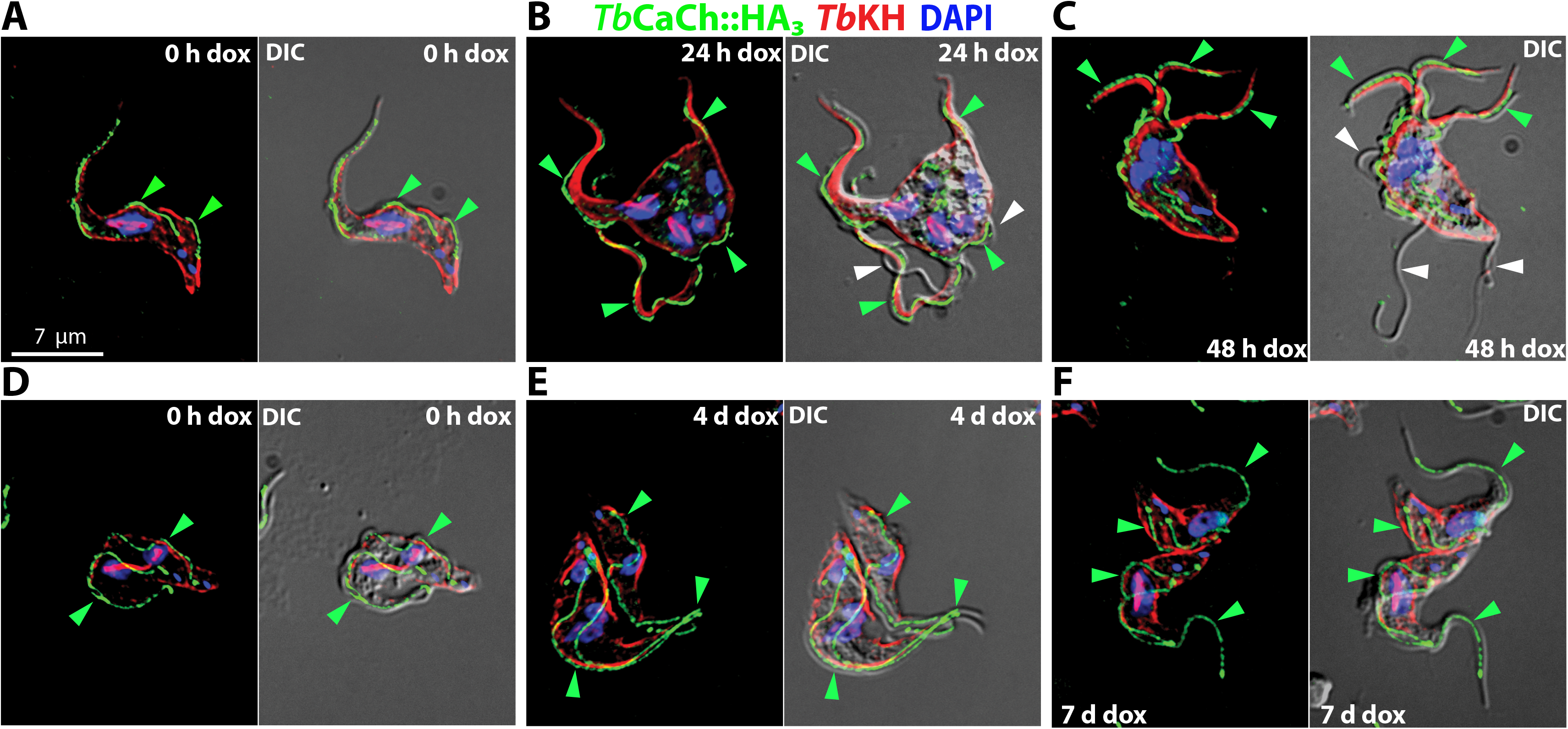
Knockdown of *Tb*KHAP1, but not *Tb*KHAP2 or *Tb*KHAP3, impairs trafficking of *Tb*CaCh::HA_3_ to the flagellum. (A-C)*Tb*CaCh::HA_3_/*Tb*KHAP1^RNAi^ BF cell line was induced with 1 μg/ml doxycycline (dox). Parasites were stained with DAPI (*blue*) and immunostained with anti-HA mAb (*Tb*CaCh::HA_3_*, green*) and anti-*Tb*KH pAb (*Tb*KH*, red*). (A) Non-induced TbCaCh::HA_3_/TbKHAP1^RNAi^ parasites (*0 h dox*), (B) parasites induced for 24 h (2*4 h dox*) and (C) parasites induced for 48 h (*48 h dox*). *Green arrowheads* in (A-C) indicate flagella where *Tb*CaCh::HA_3_ is present, and *white arrowheads* in (B-C) indicate flagella where *Tb*CaCh::HA_3_ is absent. (D-F) *Tb*CaCh::mNG/*Tb*KHAP2/3^RNAi^ BF cell line was induced with 1 μg/ml doxycycline (dox). Parasites were stained with DAPI (*blue*) and anti-*Tb*KH pAb (*Tb*KH*, red*), and mNG endogenous fluorescence was also acquired (*Tb*CaCh::mNG*, green*). Parasites were induced with doxycycline for (D) 0 days (*0 d dox*), (E) 4 days (*4d dox*), or (F) 7 days (*7d dox*). *Green arrowheads* indicate flagella where *Tb*CaCh::mNG is present. DIC images are indicated. The scale bar shown in (A) applies to all images.

In contrast, RNAi directed against *TbKHAP2/MARP-1* and *TbKHAP3/MARP-2* RNAs did not prevent trafficking of *Tb*CaCh/FS179::HA_3_ into the flagellar membrane, where it is located prior to RNAi (Fig. 7D). At both 4 d (Fig. 7E) and 7 d (Fig. 7F) post RNAi, BF parasites with multiple flagella still trafficked this channel into the flagellar membrane (green arrowheads).

### *Tb*KH-dependent trafficking of other flagellar membrane proteins

*Tb*KH is required for trafficking of *Tb*CaCh/FS179 to the flagellar membrane of BF trypanosomes, and the two protein interact with each other, as demonstrated by crosslinking pulldown assays (Sanchez et al., 2016). Is *Tb*KH important for flagellar trafficking of other membrane proteins? To address this question, we monitored the dependency of other flagellar membrane proteins on *Tb*KH for targeting to that organelle. The TrypTag project (http://tryptag.org/) has defined the subcellular location of a large number of trypanosome proteins in PF parasites (Dean et al., 2017), employing live cell microscopy of parasites expressing mNeonGreen fluorescent protein fusions, and this endeavor has identified a cohort of flagellar membrane proteins.

One such flagellar membrane protein is Tb927.7.4270, a 25 kDa protein predicted to have a N-terminal signal sequence and a single transmembrane domain (TMD) near its C-terminus. This protein is one of four paralogous proteins (Tb927.7.4230, 4260, 4270, and 4280) studied previously by Shimogawa *et al*. (Shimogawa et al., 2015) and designated the Fam79 proteins (Fam79.1, 79.2, 79.3, and 79.4, respectively). To address potential dependency upon *Tb*KH for flagellar targeting, we monitored the localization of the C-terminal mNG fusion of Tb927.7.4270/Fam97.3 in both formaldehyde fixed and live PF parasites immobilized in CyGEL (MacLean et al., 2013). As shown in Fig. 8A (0 h RNAi), this fusion protein is present in flagella and also in filaments and vesicles that emerge from the flagella, and we designate Tb927.7.4270 as extracellular vesicle membrane protein 1 or *Tb*EVMP1/Fam79.3. Multiple investigators have observed filaments and vesicles emerging from various parts of trypanosomes (Baudieri and Tomassini, 1962; Ellis et al., 1976; Molloy and Ormerod, 1964; Schepilewsky, 1912; Vickerman and Luckins, 1969; Wright and Lumsden, 1970), including the flagella, and a recent study by Szempruch *et al*. (Szempruch et al., 2016a; Szempruch et al., 2016b) has investigated such structures from BF parasites in detail and concluded that the flagellum-derived nanotubes and resulting extracellular vesicles (EVs) incorporate a cohort of parasite proteins. Furthermore, delivery of parasite-derived EVs to host red blood cells or to other trypanosomes can mediate pathogenic processes, such as erythrocyte clearance and anemia in the mammalian host or delivery of innate immune factors from a resistant to a sensitive strain of trypanosome. Hence, understanding the process for delivery of parasite proteins to these EVs is of importance for deciphering mechanisms of parasite virulence. Notably, when PF parasites expressing *Tb*EVMP1::mNG were subjected to RNAi directed against *Tb*KH for 24 h, they were strongly impaired in trafficking of this fusion protein to flagella or nanotubes (Fig. 8A, 24 h RNAi), and fluorescence often accumulated within the parasite cell body. White arrows indicate flagella that are devoid of fluorescence and which thus exhibit a trafficking defect. This result indicates that *Tb*EVMP1/Fam79.3 is dependent upon *Tb*KH, either directly or indirectly, for trafficking to the flagellum and subsequently for release into EVs. Furthermore, this trafficking defect occurs after 24 h of RNAi directed against *Tb*KH, but these PF parasites do not exhibit significant loss of viability until ~10 days of continuous RNAi (Sanchez et al., 2016), indicating that the effects of *TbKH* RNAi upon flagellar trafficking are not due to global loss of cellular functions. Notably, Fam79.1 (Tb927.7.4230) was also detected by proteomic analysis in EVs of BF trypanosomes by Szempruch *et al*. (Szempruch et al., 2016b), indicating that multiple members of this family are delivered to the membranes of EVs during the parasite life cycle.

**Fig. 8.**
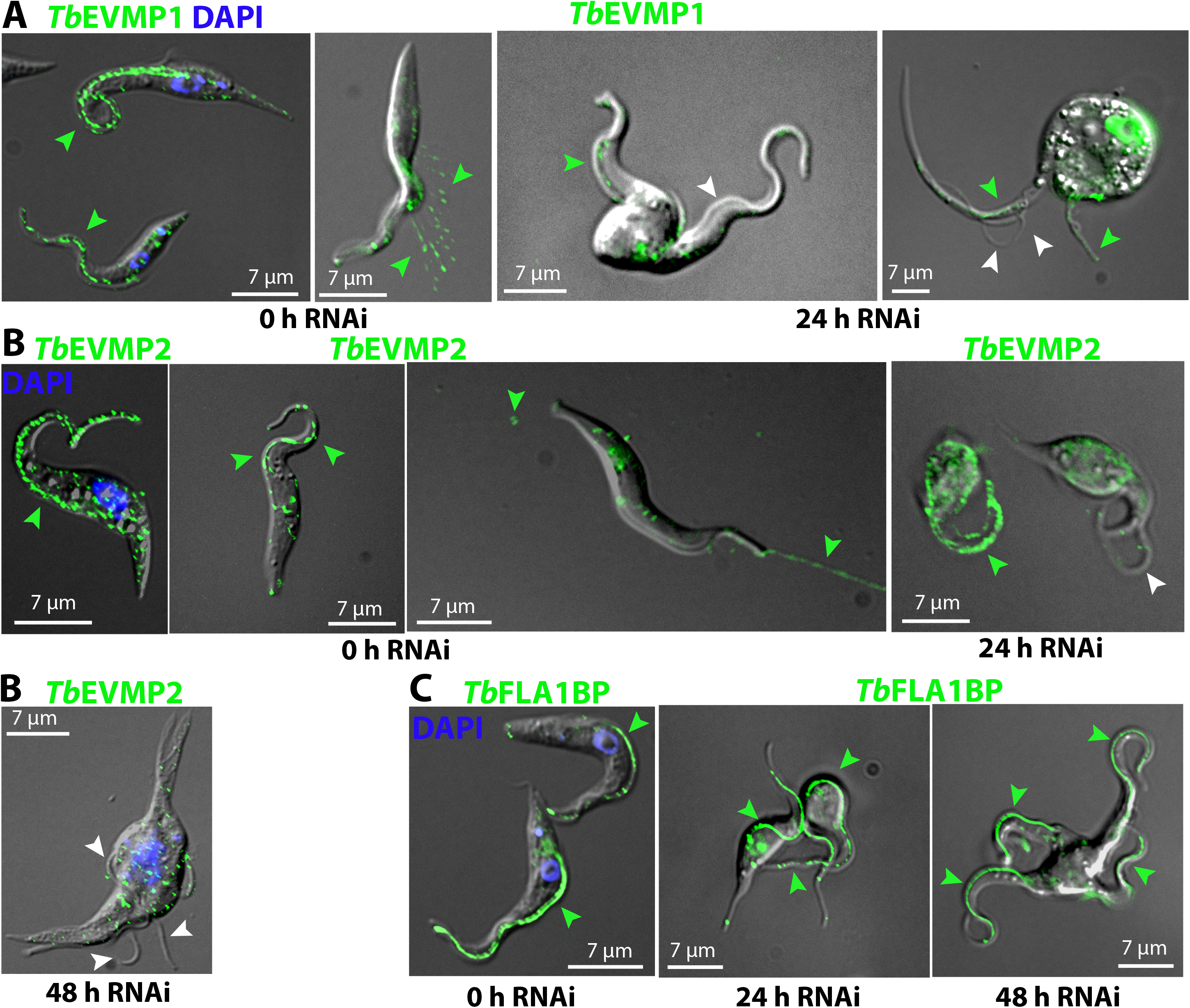
Trafficking of *Tb*EVMP1, *Tb*EVMP2, and *Tb*FLA1BP in *Tb*KH^RNAi^ BF parasites. (A) Parasites expressing *Tb*EVMP1:mNG before (*0 h RNAi*) and after (*24 h RNAi*) induction of RNAi against *TbKH* mRNA. *Green arrowheads* indicate mNG fluorescence in the FM (all panels) or in extracellular vesicles (*0 h RNAi*, right panel). *White arrowheads* indicate flagella without mNG fluorescence. (B) Parasites expressing *Tb*EVMP2::mNG before (*0 h RNAi*) and after (*24 h RNAi, 48 h RNAi*) induction of RNAi against *TbKH* mRNA. (C) Parasites expressing *Tb*FLA1BP::mNG before (*0 h RNAi*) and after (*24 h RNAi, 48 h RNAi*) induction of RNAi against *TbKH* mRNA. The left-most images in A, B, and C represent formaldehyde fixed parasites, whereas for the other images, live parasites were suspended in CyGel, which facilitates visualization of secreted extracellular vesicles.

A second flagellar membrane protein localized in TrypTag is Tb927.11.1830, designated here as *Tb*EVMP2. This 62 kDa protein has 6 predicted TMDs and is widely distributed among the kinetoplastid protists but does not have obvious orthologs outside that order, nor does it possess conserved Pfam domains (Sonnhammer et al., 1998), except for the TMDs and one predicted coiled coil. The *Tb*EVMP2::mNG fusion protein is also localized to the flagellar membrane, nanotubes, and extracellular vesicles in PF parasites (Fig. 8B, 0 h RNAi), but induction of RNAi directed against *Tb*KH also inhibits trafficking to these flagellar structures (Fig. 8B, 24 h RNAi), albeit less strongly than for *Tb*EVMP1. Although fluorescence is visible in some flagella after induction of RNAi (green arrows, righthand image for 24 h RNAI in Fig. 8B), there are some flagella that exhibit little if any fluorescence (white arrows). We designate this protein *Tb*EVMP2 and suggest that it is a second such protein that is dependent upon *Tb*KH for efficient trafficking to the surface of the flagellum.

In contrast, the mNG tagged FLA1 binding protein *Tb*FLA1BP::mNG, which is in the flagellar membrane component of the FAZ (Sun et al., 2013), traffics efficiently to the FM both before and after induction of RNAi (Fig. 8C, 0 h, 24 h, and 48 h RNAi), even in PF parasites that have multiple flagella. Hence, *Tb*FLA1BP does not require *Tb*KH for targeting to the FM, implying that there are both *Tb*KH-dependent and *Tb*KH-independent FM proteins in this parasite.

## DISCUSSION

KHARON is a cytoskeletal protein that plays multiple roles in the biology of both *T. brucei* and *L. mexicana*. In trypanosomes, *Tb*KH is critical for flagellar trafficking of *Tb*CaCh/FS179 to the FM, and failure to traffic this channel to the flagellar component of the FAZ causes flagellar detachment and arrest of cytokinesis. Nonetheless, *Tb*KH may affect cell viability by other mechanisms as well. In addition to its localization at the base of the flagellum, the protein is also present in the subpellicular cytoskeleton and mitotic spindles where it could also play roles vital to parasite viability.

In the current study, we have investigated three potential partners of *Tb*KH: *Tb*KHAP1 and the two related proteins *Tb*KHAP2/MARP1 and *Tb*KHAP3/MARP2. *Tb*KHAP1 can be crosslinked to *Tb*KH by formaldehyde indicating that the two proteins associate within 2-3 Å of each other. *Tb*KHAP2 and *Tb*KHAP3 are close enough to *Tb*KH to give a consistently positive signal using the PLA, that is within 400 Å, but the degradation of these two *Tb*KHAPs under conditions of crosslinking and solubilization has prevented us from definitively demonstrating their interaction at the near atomic level. The knockdown of *TbKHAP1* RNA has the most pronounced phenotype, strongly arresting division of BF parasites, inhibiting progression of the cleavage furrow during cytokinesis, and impairing trafficking of *Tb*CaCh/FS179 to the FM. Hence, the phenotypes of RNAi for both *TbKH* and *TbKHAP1* are similar. In contrast, efficient knockdown of *TbKHAP2/3* RNAs slows growth of BF parasites but has a much less severe effect on cell division than knockdown of *TbKHAP1* RNA. In addition, trafficking of *Tb*CaCh/FS179 to the FM is not impaired, even after 7 days of knockdown in BFs. These results imply that although both types of protein likely associate with *Tb*KH in the cytoskeleton, they play different roles. These distinctions in functions could either result from separate complexes between *Tb*KH and each partner or from different roles that each partner plays in the same complex. The association of all partners with the cytoskeleton complicates this issue, as one cannot readily separate different complexes from each other, as would be possible for cytosolic multi-protein complexes. Nonetheless, these studies confirm that *Tb*KH associates with other partner proteins that mediate its activities in different ways.

One central activity for *Tb*KH is to traffic *Tb*CaCh/FS179 to the FM, a process that is critical for integrity of the FAZ and for parasite viability. The localization of many PF proteins to their subcellular sites achieved in the TrypTag.org project (Dean et al., 2017) has identified some additional FM proteins, along with some others that were identified previously as FM components from various targeted studies (Kelly et al., 2020a), and one question of relevance is how many of these flagellar surface components rely upon *Tb*KH for organellar trafficking. For *Tb*CaCh/FS179, *Tb*KH appears to be directly involved in trafficking, since the two proteins can be crosslinked by formaldehyde and isolated as molecular partners (Sanchez et al., 2016), but it is possible that others depend upon *Tb*KH either directly or indirectly via the ability of this protein to affect various cellular processes. In this study, we have shown that two additional FM proteins, *Tb*EVMP1/Fam79.3 (Tb927.7.4270) and *Tb*EVMP2 (Tb927.11.1830) are present in both the FM and in EVs secreted from the FM in PF trypanosomes. Both proteins exhibit dependency upon *Tb*KH for trafficking to the FM, as RNAi directed against *TbKH* reduces the efficiency of their localization to this organelle. In contrast, FLA1BP, which participates in adhesion of the flagellum to the cell body by binding to the FLA1 protein in the cell body component of the FAZ (Sun et al., 2013), is not dependent upon *Tb*KH to reach the FM, confirming that both *Tb*KH-dependent and *Tb*KH-independent FM proteins exist.

EVs released from the cell body and FM of BF trypanosomes play important roles in virulence of African trypanosomes, including lysis of host erythrocytes leading to anemia, a major mechanism of trypanosome-mediated pathogenesis (Szempruch et al., 2016a; Szempruch et al., 2016b). One might anticipate that surface components of EVs could play important roles in either formation of the EV membrane or interaction of EVs with mammalian or tsetse fly tissues. *TbEVMP1* and *TbEVMP2* mRNAs are both preferentially expressed in PF trypanosomes (tritrypdb.org), but paralogs of *Tb*EVMP1, such as Tb927.7.4230 and Tb927.7.4260, are expressed at higher levels in BFs compared to PFs, suggesting potential roles for such EVMPs in both life cycle stages.

The *Tb*KH partners discovered in this study are associated primarily with the subpellicular microtubules, but there are likely to be other partners that may reside principally at the base of the flagellum or in the mitotic spindles and could be associated with distinct activities at those sites. A BioID study by Zhou *et al*. (Zhou et al., 2018) identified five spindle-associated proteins, NuSAP1, NuSAP2, Kif13-1, *Tb*MIP2, and *Tb*AUK1, that are in proximity to *Tb*KH and are thus candidates for molecular partners at the mitotic spindle. In addition, Akiyoshi and Gull (Akiyoshi and Gull, 2014) identified kinetoplast kinetochore proteins (KKTs) that associate with KHARON by CoIP/MS experiments, suggesting a possible role of KHARON in faithful chromosome segregation in kinetoplastid parasites. Molecular interaction studies of the type carried out here will be required to determine which of these proteins may be *bona fide* molecular partners with *Tb*KH and what roles KHARON complexes may be playing at the spindle. Similarly, the ability to isolate flagella with associated kinetoplast DNA and basal bodies (Oberholzer et al., 2011; Robinson and Gull, 1991; Subota et al., 2014) should facilitate identification by either BioID or TAP-MS of additional *Tb*KH partners at the base of the flagellum. Thus, it should be possible to achieve a comprehensive understanding of the role of KHARON in the cytoskeleton and the distinct functions it carries out in association with different partner proteins.

## MATERIALS AND METHODS

### Growth and transfection of *T. brucei* cell lines

BF and PF *T. brucei* cell lines were grown as described previously (Sanchez, 2013). For T7 RNA polymerase-independent driven expression the BF/pHD1313 or PF/pHD1313 clones were employed (Sanchez et al., 2016). For T7 RNA polymerase driven expression in BF, *T. brucei* 427 parasites transfected with the pSmOx (Poon et al., 2012) plasmid expressing the tetracycline repressor and T7 RNA polymerase were generated and grown in 0.1 μg/ml puromycin. For T7 RNA polymerase driven expression in PF, *T. brucei* 427 13-6 clone expressing TETR and T7 RNA polymerase was used (Wirtz et al., 1999). Linear plasmid or PCR DNA amplicons were used to transfect mid-log phase parasites as described (Sanchez et al., 2016). Transfected clones were obtained by limiting dilution according to published protocols (Burkard et al., 2007; McCulloch et al., 2004).

### Primary amino acid sequence analysis

For DNA and amino acid sequence analysis of *TbKHAPs*, ExPASy, via the SIB Bioinformatics Resource Portal (http://expasy.org), and GeneBank (http://blast.ncbi.nlm.nih.gov/Blast.cgi) or TritrypDB (http://tritrypdb.org/tritrypdb) were used.

### Inhibition of gene expression by RNAi

To inhibit the expression of *Tb*KH, *Tb*KHAPs or *Tb*FLA1 a number of cell lines were generated employing different genetic backgrounds. The BF and PF TbKH^RNAi^ and *Tb*FLA1^RNAi^ clones, for which expression is T7 polymerase-independent, were previously described (Sanchez et al., 2016), and RNAi was induced by adding doxycycline (1 μg/ml) to the culture medium. The BF and PF *Tb*KHAP1^RNAi^ clones were generated by subcloning the first 500 bp of the *TbKHAP1* (Tb927.10.1026) ORF into the pZJM RNAi vector, where expression is driven by two opposing T7 promoters (Wang et al., 2000). Similarly, BF and PF TbKHAP2/3^RNAi^ clones were generated by subcloning the last 500 bp of *TbKHAP2* (Tb927.10.10360) that is almost identical to the *TbKHAP3* (Tb927.10.10280) ORF into the pZJM RNAi vector. RNAi clones were selected by resistance to 2.5 μg/ml phleomycin and 0.1 μg/ml puromycin, and expression of dsRNA was induced by addition of 1 μg/ml doxycycline. To verify inhibition of *Tb*KHAP1-3 expression, total cell lysates were obtained from parasite cultures induced for RNAi and subjected to Western blot experiments as indicated below.

### Endogenous epitope tagging

For endogenous tagging of *Tb*KH, *Tb*KHAP1, *Tb*KHAP2, *Tb*KHAP3, CAP-15, *Tb*CaCh (FS179), *Tb*EVMP1, *Tb*EVMP2 and *Tb*FLA1BP at the N-terminus or C-terminus, different epitopes were employed as indicated in the text, following the protocol described (Dean et al., 2015). Briefly, epitope-tagging cassettes were generated by using two specific 100 nt oligonucleotides containing ~80 nt each that are homologous to the ORF region to be tagged and ~20 nt homologs to the plasmid pPOTV4 template flanking the drug resistance marker cassette, and using the universal PCR settings. Epitope tagging PCR cassettes were ethanol precipitated and resuspended in 10 μl of nucleofection buffer (Wang et al., 2000), then parasites were transfected with the purified tagging cassettes as described (Dean et al., 2015) and selected using 1.5 μg/ml G418 (15 μg ml^−1^ for PF) or 0.1 μg/ml puromycin (1 μg/ml for PF), and cloned by limiting dilution.

### Generation of Rabbit anti-*Tb*KH antibody

A custom rabbit anti-*Tb*KH polyclonal antibody (pAb) was generated by GenScript, using their 49-day antibody generation protocol. Briefly, two rabbits were injected with 200 μg of His_6_::*Tb*KH, representing amino acids 43 - 411, emulsified in Freund’s complete adjuvant. The rabbit was boosted 3 times at 14-day intervals with 200 μg of His_6_::*Tb*KH emulsified in Freund’s incomplete adjuvant. Antibody specificity for *Tb*KH was evaluated by western blot comparing the reactivities of the rabbit serum from immunized rabbits to *T. brucei* protein lysates from wild-type cells and N-BirA*::*Tb*KH cell line (Fig 3A). Proteins were immunodetected using 1:2500 dilution of the rabbit anti-*Tb*KH polyclonal antibody and 1:15,000 dilution of goat anti-rabbit-HRP antibody (Sigma-Aldrich). The chemiluminescent protocol was used for developing as indicated below.

### Immunofluorescence microscopy

For immunofluorescence microscopy, 5 × 10^6^ parasites were centrifuged at 1000 X g for 5 min and washed twice at room temperature with phosphate buffered saline pH 7.2 (PBS) containing 10 mM glucose. The cell pellet was resuspended in 4% paraformaldehyde in PBS, pH 7.2 and incubated for 15 min at room temperature, cells were centrifuged as descried above and washed once with PBS, resuspended in 100 μl PBS, spotted onto poly-L-lysine coated coverslips, permeabilized with 0.1% Triton X-100 in PBS for t min, and washed 3X with PBS. Then parasites were blocked with 2% goat serum, 0.01 sodium azide, 0.01 saponin in PBS (blocking solution) for 1 h at room temperature, rinsed 3 X with PBS, and incubated with primary antibodies for 1 h at room temperature. The following primary antibodies were employed: 1:250 dilution rabbit anti-*Tb*KH pAb (reported in this work), 1:500 dilution mouse anti-HA monoclonal antibody (mAb) (BioLegend, Cat # MMS-101R), and 1:1000 dilution mouse anti-α-tubulin mAb (Sigma-Aldrich, Cat. # T5168). Subsequently, cells were rinsed as before and incubated with a 1:1000 dilution of secondary antibodies coupled to Alexa Fluor dyes (Molecular Probes) as follows: Alexa Fluor® 488 goat anti-mouse IgG (H+L) (Cat. # A11001), Alexa Fluor® 594 goat anti-mouse IgG (H+L) (Cat. # A11005), Alexa Fluor® 594 goat anti-rabbit IgG (H+L) (Cat. # A11012) and Alexa Fluor® 488 goat anti-rabbit IgG (H+L) (Cat. # A11008), as indicated, in blocking solution for 1h at room temperature in the dark. Cover slips were rinsed 3 X with PBS and mounted onto slides using DAPI Fluoromount-G (SouthernBiotech). Fluorescence images were obtained using a wide field deconvolution system (Applied Precision Instruments, Inc.) consisting of an inverted Nikon TE 200 Eclipse microscope, a Kodak CH350 CCD camera, and the Deltavision operating system. Images were acquired using a 60 X objective and 1.25 X magnification in a 1024 × 1024 format, and deconvolved using SoftWoRx software. Adobe Photoshop CC and Adobe Illustrator CC (Adobe Systems Inc.) were used to create image compositions.

### Flagellar protein purification, formaldehyde crosslinking, pull down, and Western blot assay

Flagellar purification was performed from BF parasites (Fig. 3A,B and Fig. S1A,B) as described (Subota et al., 2014) with slight changes. Briefly, BF *Tb*KH1^RNAi^, *Tb*FLA1^RNAi^, *Tb*KHAP1^RNAi^ and *Tb*KHAP2/3^RNAi^ clones were grown in 1 μg/ml doxycycline for the indicated times and pelleted at 420 X g for 10 min. Cell pellets were washed with buffer A (25 mM Na^+^-tricine, pH 7, 1% BSA, 0.1 mM CaCl_2_, 0.2 mM EDTA, 5 mM MgCl_2_ and 12 mM β-mercaptoethanol) containing 0.32 M sucrose and centrifuged at 420 X g for 10 min. Cell pellets were gently resuspended at 3 × 10^8^ parasites/ml in buffer A plus 0.3 M sucrose, transferred into Eppendorf tubes and vortexed for 5 min or until microscopic verification of flagellum detachment, followed by centrifugation at 420 X g for 10 min. Supernatants were recovered and centrifuged at 16,000 X g for 20 min at 4 °C. Pellets containing the isolated flagella were resuspended in 200 μl of PBS and used for immunofluorescence microscopy. Also, flagellar purification from PF clones without induction of RNAi (Fig. 3C and Fig. S1C) was performed by isolation of cytoskeletons followed by treatment with 1 mM CaCl_2_, according to Imhof et al. (Imhof et al., 2019).

For formaldehyde crosslinking and pull down, cell lines were prepared that expressed *Tb*KH1 endogenously tagged with the His_10_ epitope at the N-terminus (His_10_::*Tb*KH1) and also co-expressed with a protein of interest endogenously tagged at its N- or C-terminus with a HA_3_ or V5_3_ epitope (HA_3_::*Tb*KHAP1, HA_3_::*Tb*KHAP2, *Tb*KHAP2::HA_3,_ V5_3_::*Tb*KHAP3, HA_3_::*Tb*KHAP3 and *Tb*CAP15::HA_3_). Parasites were washed once with PBS and pelleted at 1000 X g for 10 min, and cell pellets resuspended in 9.37 ml PBS plus 0.63 ml 16% formaldehyde-EM grade (Polysciences, Inc) and incubated at room temperature for 10 min. For non-crosslinked control samples, PBS was added instead of formaldehyde. Subsequently, 1 ml of 2.5 M glycine in PBS was added to the crosslinked samples and incubated at room temperature for 5 min, crosslinked parasites were pelleted at 1000 X g for 10 min and washed twice with PBS. Then, cell pellets with or without crosslinking were resuspended in 1 ml of Buffer 1 (8 M urea, 300 mM NaCl, 0.5% Nonidet P-40, 50 mM NaH_2_PO_4_, 50 mM Tris, pH 7.0, 10 mM imidazole) on ice. Samples were sonicated 3 times on ice at 50% max amplitude (Sonic Dismembrator, 500W, Fisher Scientific) for 10 s with 30 s between pulses. A 2.5% aliquot of this protein lysate was saved as the protein lysate fraction (LF).

The remainder of the protein was incubated with 50 μl of Ni-NTA Magnetic Beads (New England Biolabs, Inc.) on a rocker for 45 min at room temperature. The beads were washed three times according to the manufacturer’s instructions using Buffer 1 plus 20 mM imidazole. Finally, bound protein complexes were eluted with 100 μl Buffer 1 containing 500 mM imidazole. Crosslinking was reversed by boiling protein samples for 30 min in 1X Bolt™-LDS sample buffer (Life Technologies) containing 10 mM DTT.

Protein extracts were prepared and analyzed by Western blot employing Bolt^™^ 4-12% Bis-Tris Mini Protein Gels or NuPage™ 3-8% Tris-acetate gels, Mini Gel Tank and Mini Blot Module following the manufacturer’s instructions (Life Technologies). Proteins were transferred onto PVDF membranes (Millipore). Protein immunodetection was done using rabbit anti-*Tb*KH pAb at 1:2500 dilution, mouse anti-HA mAb (BioLegend, Cat # MMS-101R) 1:2500 dilution, mouse anti-V5 mAb (Invitrogen, Cat # MA5-15253) 1:2500 dilution, and mouse anti-α tubulin mAb (Sigma-Aldrich, Cat. # T5168) 1:10,000 dilution. Goat anti-rabbit-HRP (Sigma-Aldrich) 1:15,000 dilution and goat anti-mouse-HRP (Jackson ImmunoResearch Laboratories, Cat. # 115-03-174, Lot # 117119) were used as secondary antibodies and Western blots were developed using the SuperSignal^™^ West Pico Plus Chemiluminescent Substrate (Thermo Fisher Scientific) and an Image Quant LAS 400 (GE Healthcare) scanner was employed to acquire luminescent images. Adobe Photoshop CC and Adobe Illustrator CC (Adobe Systems Inc.) were used to create image compositions.

### Proximity ligation assay (PLA)

Mid-log BF or PF parasites were harvested by centrifugation at 1000 x g for 10 min, washed once in PBS and fixed with 4% paraformaldehyde in PBS for 15 min at room temperature. Fixed cells were attached to cover slips and permeabilized as indicated above for immunofluorescence analysis. Subsequently the PLA protocol was followed according to the Duolink In Situ Red Starter Kit Mouse/Rabbit (Millipore Sigma) instructions. Briefly, after blocking cells were incubated with 1:250 dilution of rabbit anti-*Tb*KH pAb and 1:500 dilution of mouse anti-HA mAb (BioLegend, Cat# MMS-101R). As a negative control, cells were incubated only with 1:250 dilution of the rabbit anti-*Tb*KH pAb. Then PLA species-specific secondary antibodies with minus and plus oligonucleotide probes were added, followed by ligation, amplification and hybridization with specific red-fluorescent oligonucleotides to allow detection by fluorescence microscopy. Samples were mounted and imaged as described for immunofluorescence analysis.

## Acknowledgements

We appreciate the expert advice and support of the staff of the Advanced Light Microscopy Core in the Jungers Center for Neurosciences at Oregon Health & Science University. We acknowledge discussions with Dr. Samuel Dean (University of Oxford) concerning new flagellar membrane proteins identified by the TrypTag project.

## Author contributions

Conceptualization: M.A.S. and S.M.L.; Data collection: M.A.S.; Data analysis: M.A.S. and S.M.L.; Writing and editing: M.A.S. and S.M.L.; Funding acquisition and project administration: S.M.L.

