## Supplemental Information for "The *Trypanosoma brucei* Cytoskeletal Protein KHARON Associates with Partner Proteins to Mediate Both Cytokinesis and Trafficking of Flagellar Membrane Proteins"

Figure S1

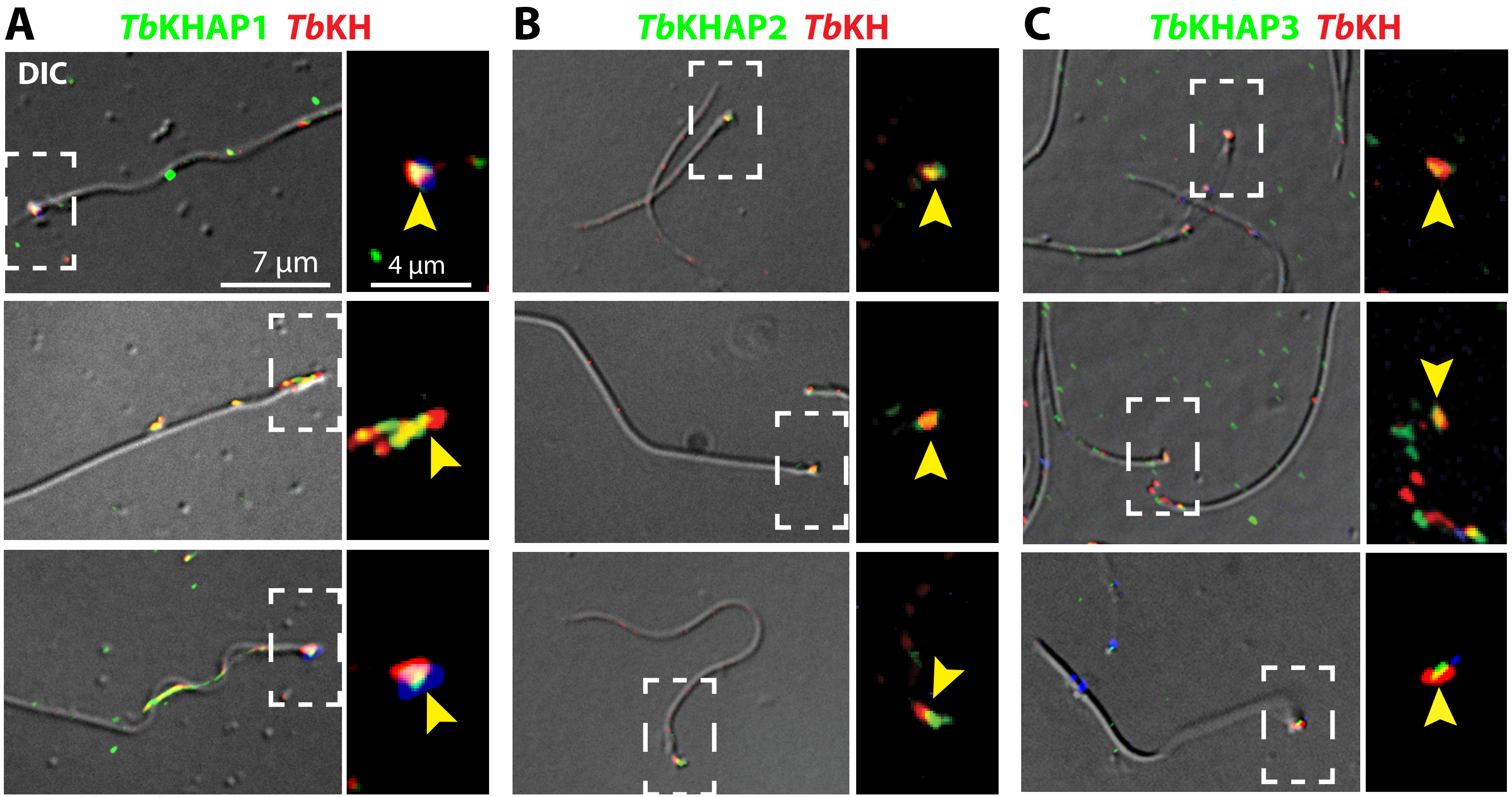


**Fig. S1.** Images of 3 isolated flagella each, showing overlap of TbKH (*red*) with A) $HA_3$::TbKHAP1, B) $HA_3$::TbKHAP2, and C) $HA_3$::TbKHAP3 (*green*). Conditions for preparation of flagella are the same as in Fig. 3. For each panel, the images at the left are DIC plus fluorescence, with the scale bar representing 7 µm in all such images, whereas the images at the right are expanded views of the region within the dashed white box shown as fluorescence images only, and with the single scale bar representing 4 µm in all such images. The *yellow arrowhead* points to overlap between the red and green fluorescence.

Figure S2

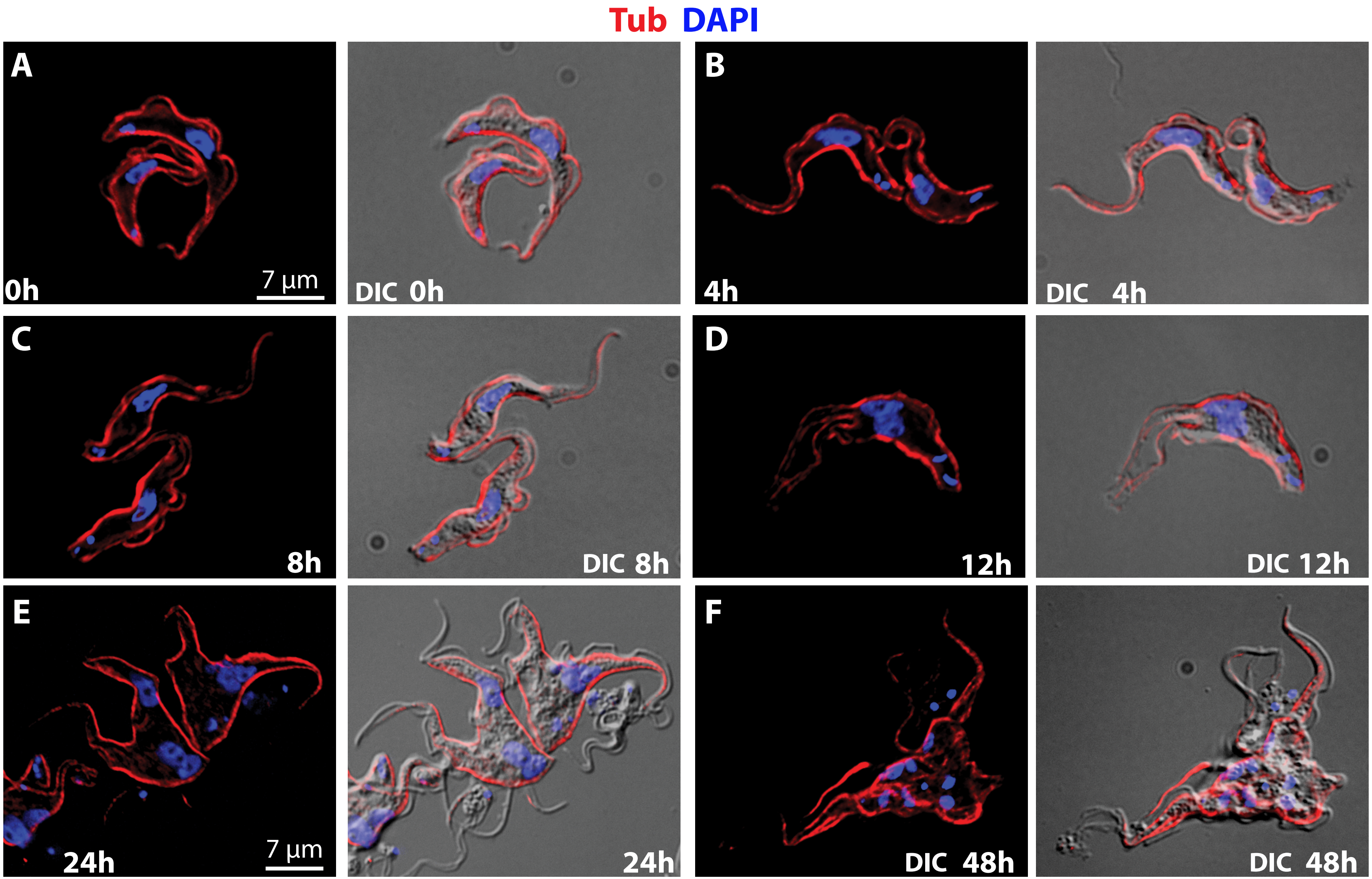


**Fig. S2.** Images of BF parasites subjected to RNAi against *TbKHAP1* mRNA for increasing periods of time. Images were recorded as in Fig. 5E,F, with *red* representing α-tubulin (Tub) and *blue* DAPI staining. The times indicated in each panel are the number of h after addition of doxycycline to induce RNAi. The 7 µm scale bar in A refers to images in A-D, and the scale bar in E refers to images in E-F. For each panel, images at the left are fluorescence and those at the right represent overlap of fluorescence with DIC.
